# DMT alters cortical travelling waves

**DOI:** 10.1101/2020.05.06.080937

**Authors:** Andrea Alamia, Christopher Timmermann, Rufin VanRullen, Robin L. Carhart-Harris

**Author notes:** These authors are first-coauthors (Andrea Alamia, Christopher Timmermann). These authors share equal senior contribution (Rufin VanRullen, Robin L. Carhart-Harris).

## Abstract

Psychedelic drugs are potent modulators of conscious states and therefore powerful tools for investigating their neurobiology. N,N, Dimethyltryptamine (DMT) is a particularly interesting serotonergic psychedelic that can rapidly induce an extremely immersive state of consciousness characterized by vivid and elaborate visual imagery. In the present study, we investigated the electrophysiological correlates of the DMT-induced altered state, by recording EEG signals from a pool of participants receiving DMT and (separately) placebo (saline), intravenously, while instructed to keep their eyes closed (i.e. ‘resting state’). Consistent with our prior hypotheses, results revealed a spatio-temporal pattern of cortical activation (i.e., travelling waves) similar to that elicited by visual stimulation. Moreover, the typical top-down alpha-band rhythms of closed-eyes rest (i.e. a backward travelling wave) were significantly decreased, while the bottom-up ‘forward travelling wave’, was significantly increased. These results support a recent model proposing that psychedelics reduce the ‘precision-weighting of priors’, thus altering the balance of top-down versus bottom-up information passing, where properties of backward waves are considered correlates of this precision weighting. The robust hypothesis-confirming nature of the present findings imply the discovery of an important mechanistic principle underpinning psychedelic-induced altered states – i.e. reduced backward and increased forward travelling waves - and lend further support to prior assumptions about the functional significance of cortical travelling waves.

## Introduction

N,N, Dimethyltryptamine (DMT) is a mixed serotonin receptor agonist that occurs endogenously in several organisms ^1,2^ including humans ^3^. DMT, which is a classic psychedelic drug, is also taken exogenously by humans to alter the quality of their consciousness. For example, synthesized compound is smoked or injected but it has also been used more traditionally in ceremonial contexts (e.g. in Amerindian rituals). When ingested orally, DMT is metabolized in the gastrointestinal (GI) system before reaching the brain. Its consumption has most traditionally occurred via drinking ‘Ayahuasca’, a brew composed of plant-based DMT and *β* –carbolines (monoamine oxidize inhibitors), which inhibit the GI breakdown of the DMT ^4^. Modern scientific research has mostly focused on intravenously injected DMT. Administered in this way, DMT’s subjective effects have a rapid onset, reaching peak intensity after about 2-5 minutes and subsiding thereafter, with negligible effects felt after about 30 minutes ^5–7^.

Previous electrophysiological studies investigating changes in spontaneous (resting state) brain function elicited by ayahuasca have reported consistent broadband decreases in oscillatory power ^7,8^, while others have noted that the most marked decreases occur in α-band oscillations (8-12 Hz) ^9^. Alpha decreases correlated inversely with the intensity of ayahuasca-induced visual hallucinations ^10^ and are arguably the most reliable neurophysiological signature of the psychedelic state identified to-date ^11^ – with increased signal diversity or entropy being another particularly reliable biomarker ^12^. In the first EEG study of the effects of pure DMT on on-going brain activity, marked decrease in the α and β (13 – 30 Hz) band power were observed as well as increases in signal diversity ^7^. Increases in lower frequency band power (δ = 0.5 – 4 Hz and θ = 4 – 7 Hz) also became evident when the signal was decomposed into its oscillatory component. Decreased alpha power and increased signal diversity correlated most strongly with DMT’s subjective effects – consolidating the view that these are principal signatures of the DMT state, if not the psychedelic state more broadly.

Focusing attention onto normal brain function, outside of the context of psychoactive drugs, electrophysiological recordings in cortical regions reveal distinct spatio-temporal dynamics during visual perception, which differ considerably from those observed during closed-eyes restfulness. It is possible to describe these dynamics as oscillatory ‘travelling waves’, i.e. fronts of rhythmic activity which propagate across regions in the cortical visual hierarchy ^13–15^. Recent results showed that travelling waves can spread from occipital to frontal regions during visual perception, reflecting the forward bottom-up flow of information from lower to higher regions. Conversely, top-down propagation from higher to lower regions appears to predominate during quiet restfulness^16–18^.

Taken together these results compel us to ask how travelling waves may be affected by DMT, particularly given their association with predictive coding ^19^ and a recent predictive coding inspired hypothesis on the action of psychedelics – which posits decreased top-down processing and increased bottom up signal passing under these compounds ^20^. Moreover, DMT lends itself particularly well to the testing of this hypothesis as its visual effects are so pronounced. Given that visual perception is associated with an increasing in forward travelling waves and eyes-closed visual imagery under DMT can feel as if one is ‘seeing with eyes shut’ – does a consistent increase in forward travelling waves under DMT account for this phenomenon?

Here we sought to address these questions by quantifying the amount and direction of travelling waves in a sample of healthy participants who received DMT intravenously, during eyes-closed conditions. We hypothesized that DMT acts by disrupting the normal physiological balance between top-down and bottom-up information flow, in favor of the latter ^20^. Moreover, does this effect correlate with the vivid ‘visionary’ component of the DMT experience? Providing evidence in favor of this hypothesis would indicate that forward travelling waves do play a crucial role in conscious visual experience, irrespective of the presence of actual photic stimulation.

## Results

### Quantifying travelling waves

We measure the waves’ amount and direction with a method devised in our previous studies^16,17^. We slide a one-second time-window over the EEG signals (with 0.5s overlap). For each time window, we generate a 2D map (time/electrodes) by stacking the signals from 5 central mid-line electrodes (Oz to FCz, see figure 1). For each map, we then compute a 2D-FFT, in which the upper- and lower-left quadrant represent the power of forward (FW) and backward (BW) travelling waves, respectively (since the 2D-FFT is symmetrical around the origin, the lower- and upper-right quadrants contain the same information). From both quadrants we extracted the maximum values, representing the raw amount of FW and BW waves in that time-window. Next, we performed the same procedure after having shuffled the electrodes’ order, thereby disrupting spatial information (including the waves’ directionality) while retaining the same overall spectral power. In other words, the surrogate measures reflect the amount of waves expected solely due to the temporal fluctuations of the signal. After having computed the maximum values for the FW and BW waves of the surrogate 2D-FFT spectra one hundred times (and averaging the 100 values), we compute the net amount of FW and BW waves in decibel (dB), by applying the following formula:

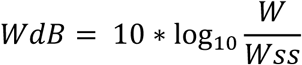

where W represents the maximum value extracted for each quadrant (i.e. forward FW or backward BW), and Wss the respective surrogate value.

**Figure 1:**
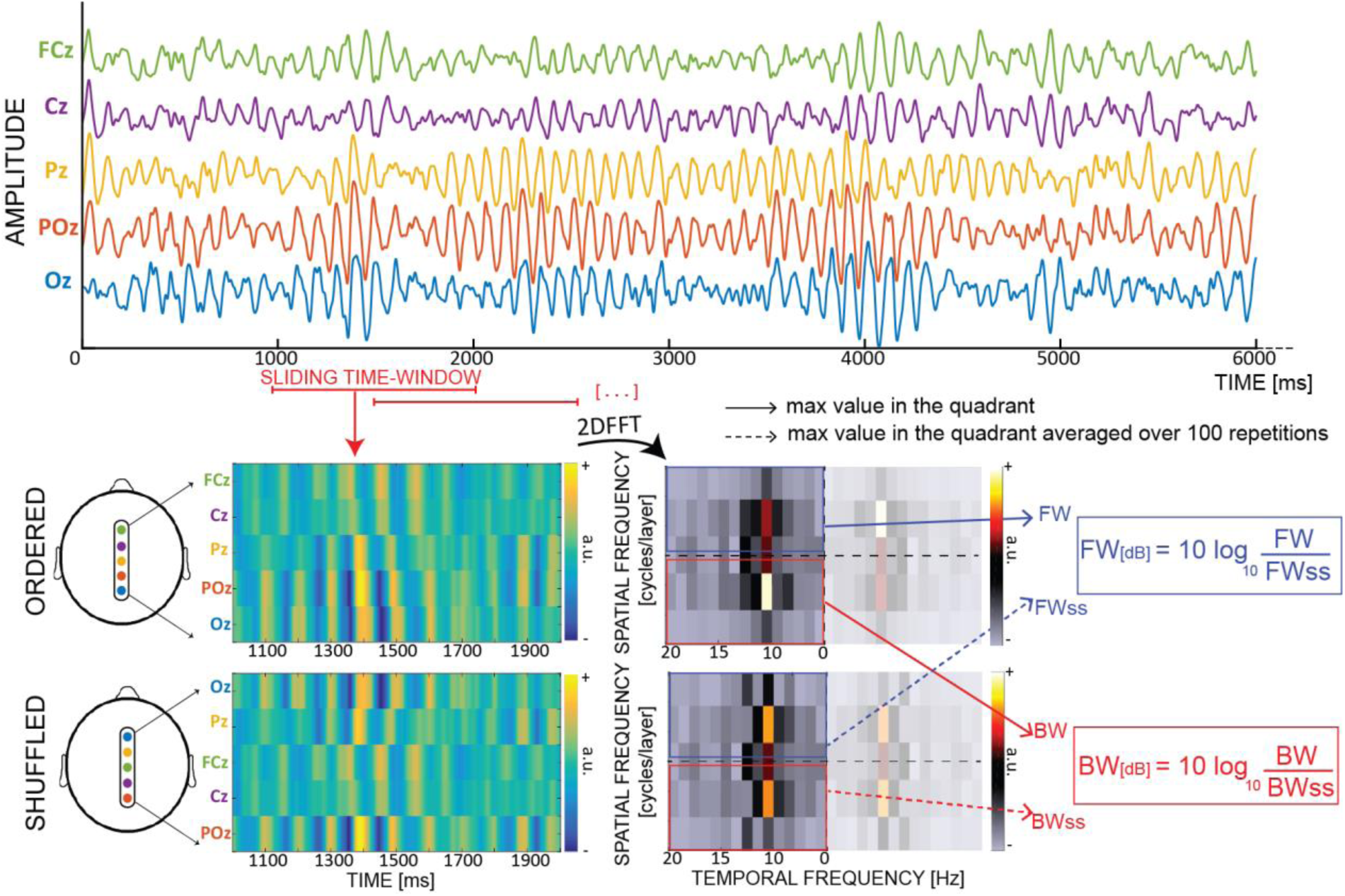
Quantifying cortical waves. From each 1-s EEG epoch we extract a 2D-map, obtained by stacking signals from 5 midline electrodes. For each map we compute a 2D-FFT in which the maximum values in the upper- and lower-left quadrants represent respectively the amount of forward (FW – in blue) and backward (BW – in red) waves. For each map, we also compute surrogate values by shuffling the electrodes’ order 100 times, so as to retain temporal fluctuations while disrupting the spatial structure of the signals (including any travelling waves). Eventually, we compute the wave strength in decibel (dB) by combining the real and the surrogate values.

Importantly, this value-expressed in decibel-represents the net amount of waves against the null distribution. In other words, it is informative to compare this value to zero, to assess the significance of waves. On the other hand, a direct comparison between FW and BW waves in each time-bin is not readily interpretable, as it is possible to simultaneously record waves propagating in both directions—as observed during visual stimulation epochs (see below).

### Does DMT influence travelling waves?

After defining our measure to assess the waves’ amount and direction, we investigated whether the intake of DMT alters the cortical pattern of travelling waves. Participants underwent two sessions in which they were injected with either placebo or DMT (see Methods for details). Importantly, during all the experiments they rested in a semi-supine position, with their eyes closed. EEG recordings were collected 5 minutes prior to drugs administration and up to 20 minutes after. The left column of figure 2A shows the amount of BW and FW waves in the 5 minutes preceding and following drug injection (either placebo or DMT). As previously reported ^16^, during quiet closed-eyes restfulness a significant amount of BW waves spread from higher to lower regions (as confirmed by a Bayesian t-test against zero for both DMT and Placebo conditions, BFs_10_>>100), whereas no significant waves propagate in the opposite FW direction (Bayesian t-test against zero: BFs_10_<0.15). However, after DMT injection the cortical pattern changed drastically: the amount of BW waves decreased (but remaining significantly above zero - BFs_10_=12.6) whereas the amount of FW waves increased significantly above zero (BF_10_=5.4). All in all, these results obtained by comparing the amount of waves before and after injection (pre-post factor) of Placebo or DMT (drug factor) were confirmed by two Bayesian ANOVA performed separately on BW and FW waves (all factors including interactions reported BFs_10_ >> 100). Additionally, we ran a more temporally precise analysis, on a minute by minute scale, testing the amount of FW and BW waves in the two conditions, as shown in the right panels of figure 2A. In line with previous studies ^5,7,21^, the changes in cortical dynamics appeared rapidly after intravenous DMT injection, and faded off within dozens of minutes. Confirming our previous analysis, we observed an increase in FW waves (asterisks in the lower right panel of figure 2A show FDR-corrected significant p-values when testing against zero) and a decrease in BW waves, which nonetheless remained above zero (all FDR-corrected p-values<0.05). To our initial surprise, the dynamics elicited by DMT injection were remarkably reminiscent of those observed in another study, in which healthy participants alternated visual stimulation with periods of blank screen, without any drug manipulation^17^. Although a direct comparison is not statistically possible (because the two studies involved distinct subject groups and different EEG recording setups), we indirectly investigated the similarities between these two scenarios.

**Figure 2:**
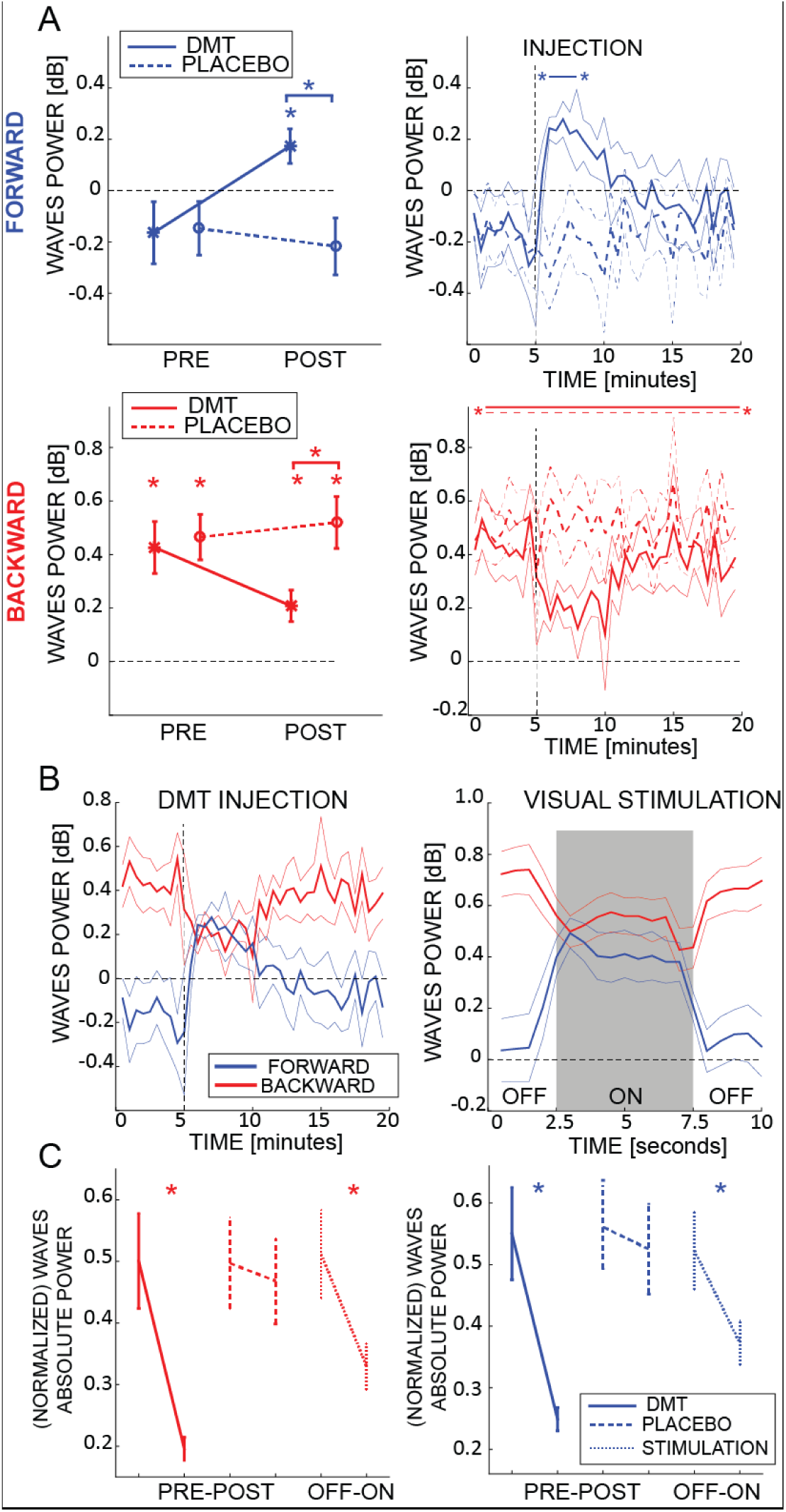
DMT influences cortical travelling waves. A) In the left panels the net amount of FW (blue, upper panel) and BW (red, lower panel) waves is represented pre- and post-DMT infusion. While BW waves are always present, FW waves only rise significantly above zero after DMT injection, despite participants having closed eyes. Asterisks denote values significantly different than zero, or between conditions. The panels to the right describe the minute-by-minute changes in the net amount of waves. Asterisks denote FDR-corrected p-values for amount of waves significantly different than zero. B) Comparison between the waves’ temporal evolution after DMT injection (left panel) and with or without visual stimulation (right panel, from a different experiment in which participants, with open eyes, either watched a visual stimulus or a blank screen^17^). Remarkably, the waves’ temporal profiles are very similar in the two conditions, for both FW and BW. C) Comparison between changes in absolute power (as extracted from the 2D-FFT, i.e. FW and BW in figure 1) due to DMT, placebo and visual stimulation. Remarkably, true photic visual stimulation and eyes-closed DMT induce comparably large reductions in absolute power. In fact, the effect with DMT appears to be even more pronounced (formal contrast not appropriate). Note that in the previous panels the changes in the net amount of waves were reported in dB, and occurred irrespective of the global power changes measured in panel C.

### Comparison with perceptual stimulation

We recently showed that FW travelling waves increase during visual stimulation, whereas BW waves decrease, in line with their putative functional role in information transmission ^17^. In figure 2B, for the sake of comparison, we contrast the cortical dynamics induced by DMT (left panel) with the results of our previous study (right panel^17^), in which participants perceived a visual stimulus (label ‘ON’) or stared at a dark screen (label ‘OFF’). Remarkably, mutatis mutandis, both FW and BW waves share a similar profile across the two conditions, increasing and decreasing respectively following DMT injection or visual stimulation. If we consider the absolute (maximum) power values derived from the 2D-FFT of each map (i.e., before estimating the surrogates and the waves’ net amount in decibel) as an estimate of spectral power, we can read the results reported in figure 2C as an overall decrease in oscillatory power following DMT injection, more specifically in the frequency band with the highest power values (i.e. alpha band, but see next paragraph) ^7–9,11^. Such decrease in oscillatory power is also matched by a similar decrease induced by visual stimulation (all Bayesian t-test BFs_10_>>100). These results demonstrate that, despite participants having their eyes-closed throughout, DMT produces spatio-temporal dynamics similar to those elicited by true visual stimulation. These results therefore shed light on the neural mechanisms involved in DMT-induced visionary phenomena.

### Does DMT influence the frequency of travelling waves?

Previous studies showed that DMT alters specific frequency bands (e.g. alpha-band^9^), mostly by decreasing overall oscillatory power ^7,8^. Here, we investigated whether DMT influences not only the waves’ direction but also their frequency spectrum. We compared the frequencies of the maximum peaks extracted from the 2D-FFT (see figure 1) before and after DMT or Placebo injection. Before infusion, both FW and BW waves had a strong amount of waves oscillating in the alpha range (figure 3, left panel). Remarkably, following DMT injection, the waves’ spectrum changed drastically, with a significant reduction in the alpha-band, coupled with an increase in the delta and theta bands, for both FW and BW waves (all BFs_10_>100). This result corroborates a previous analysis performed on EEG recordings from the same dataset ^7^ as well as independent data pertaining to O-Phosphoryl-4-hydroxy-N,N-DMT (psilocybin), a related compound ^11^.

**Figure 3:**
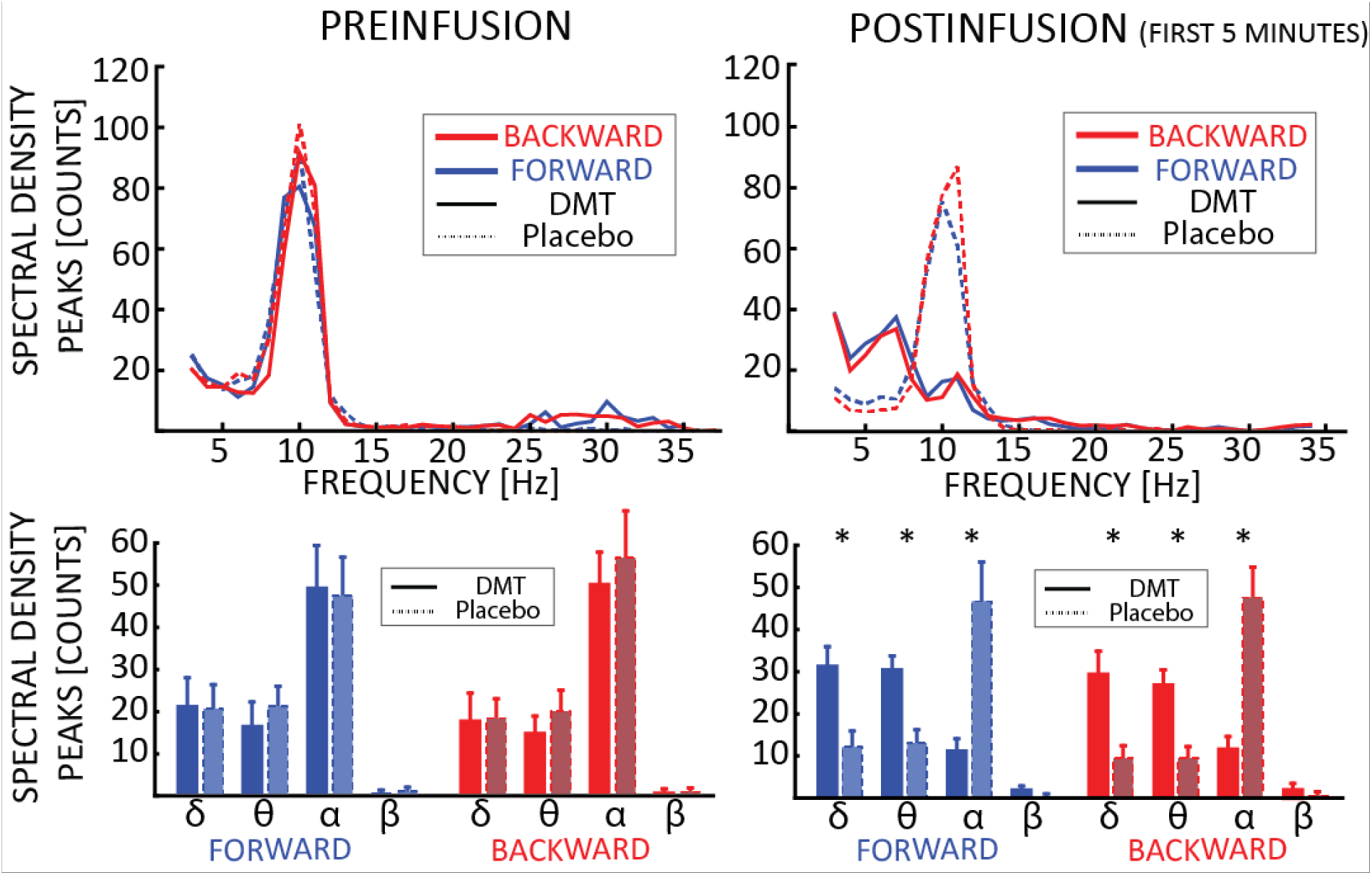
DMT influences the frequency of the travelling waves. Upper panels show the spectra for BW and FW waves pre-(left) and post-(right) infusion. The waves’ frequencies are computed from the maximum value from each quadrant in the 2D-FFT map; the histogram reflects the number of 1s time-windows having a wave peak at the corresponding frequency. The lower panels report the average between participants for each frequency band (1<δ<4Hz, 4<θ<7Hz, 8<α<12Hz, 13<β<30Hz). Notably, DMT significantly reduces α and β band oscillations, while enhancing δ and θ. Asterisks denote significant differences between DMT and Placebo conditions.

### What’s the relationship between FW and BW waves?

From the left panel of figure 2B, it seems that during the first minutes after DMT injection, both FW and BW waves are simultaneously present in the brain. In an attempt to understand the overall relationship between FW and BW waves, we focused on the minutes when both BW and FW waves were significantly larger than 0 (minutes 2 to 5 after DMT injection, see figure 2A). On these data we performed a moment-by-moment correlation between their respective net amount (as measured in decibel – see figure 1). We found a clear and significant negative relationship, very consistent across participants and irrespective of DMT injection (difference between Pre and Post, Bayesian t-test BF_10_ = 0.225; figure 4, first panel). This result demonstrates that, in general, FW waves tend to be weaker whenever BW waves are stronger, and vice-versa. In other words, FW and BW remain present after drug injection, sum to a consistent total amount, and remain inversely related; it is only the ratio of contribution from each that changes after DMT (i.e. less BW, more FW waves).

**Figure 4:**
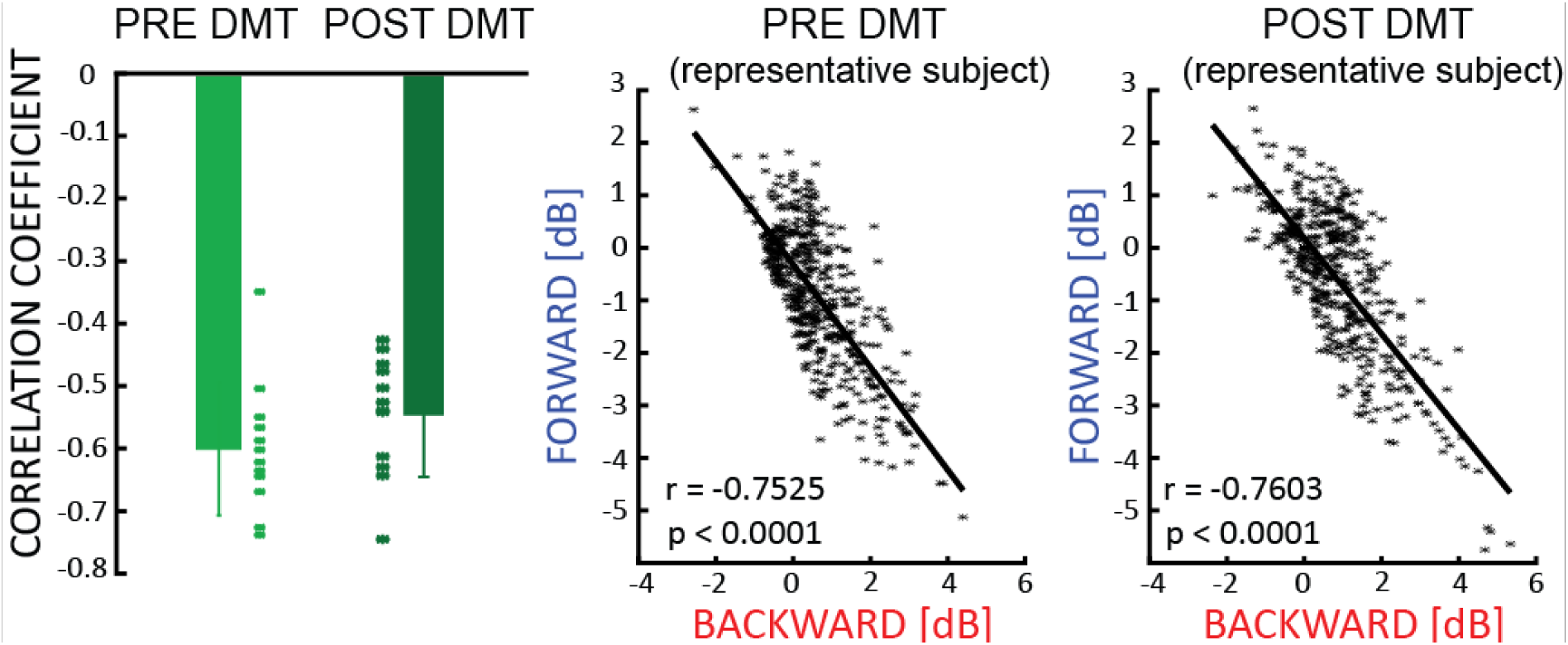
Travelling waves directions. There is a negative correlation between the net amount of FW and BW waves, which is not influenced by the ingestion of DMT (left panel). The middle and the right panel show the relationship for a typical subject pre- and post-DMT injection.

### Is there a correlation between waves and subjective reports?

We investigated whether changes in travelling waves under DMT correlated with the subjective effects of the drug. Specifically, for 20 minutes after DMT injection participants provided an intensity rating every minute and, when subjective effects faded, participants filled various questionnaires that addressed different aspects of the experience (see Timmermann et al., 2019 for details). First, we found a robust correlation between minute-by-minute intensity rates and the amplitude of the waves, as shown in the first panel of figure 5. This result reveals that the developing intensity of the drug’s subjective effects and changes in the amplitude of waves correlate positively (FW) or negatively (BW) across time, both peaking a few minutes after drug injection. Second, treating each time point independently, we again correlated intensity ratings with the amount of each wave type, across subjects. The middle panel of figure 5 shows a clear trend for the correlation coefficients over time. Despite the limited number of data-points (n=12), the correlation coefficients reach high values (∼0.4), implying that, around the moment where the drug had its maximal effect (2-5 minutes after injection), those subjects who reported the most intense effects were also those who had the strongest travelling waves in the FW direction, and the weakest waves in the BW direction. Finally, we correlated the amount of FW and BW waves with ratings focused specifically on visual imagery: remarkably, all the correlations between each questionnaire item correlated positively with the increased amount of FW waves under DMT. As the same relationship was not apparent for the BW waves, this consolidates the view that visionary experiences under DMT correspond to higher amounts of FW waves in particular. Taken together with previous results from visual stimulation experiments independent of DMT^17^, these data strongly support the principle cortical travelling waves (and increased FW waves in particular) correlate with the conscious visual experiences, whether induced exogenously (via direct visual stimulation) or endogenously (visionary or hallucinatory experiences).

**Figure 5:**
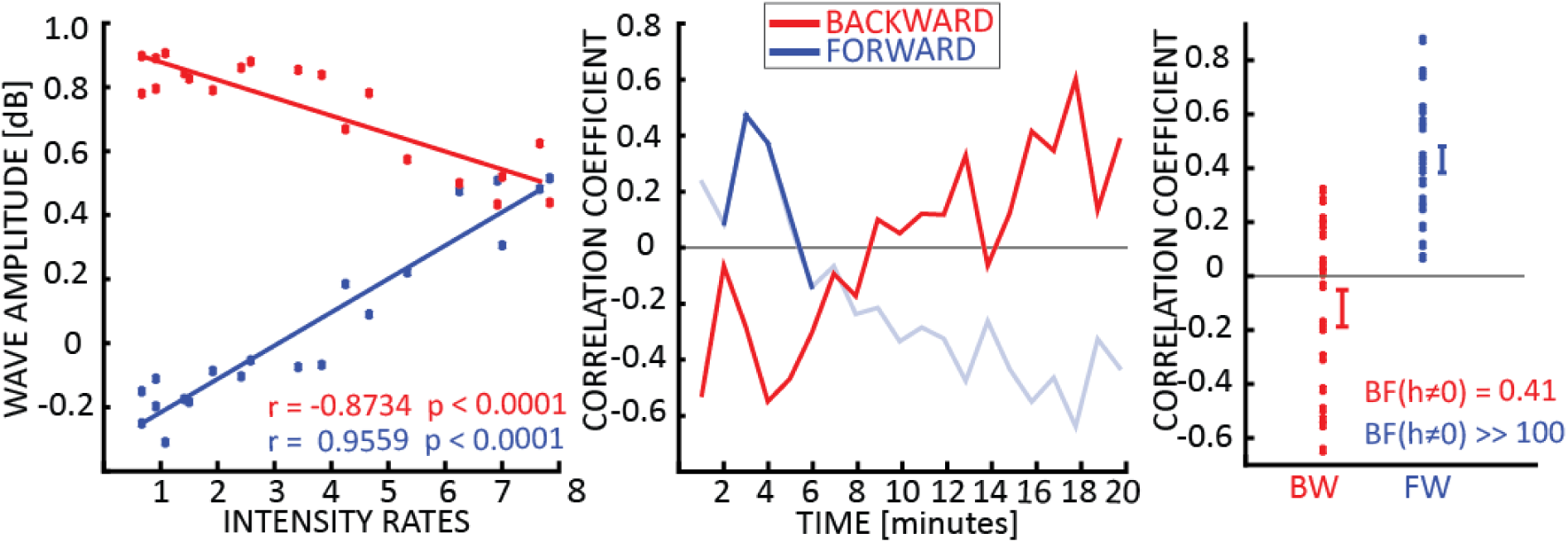
Travelling waves vs subjective ratings. The first panel shows the correlation between intensity rate and waves amplitude across time-points. Each dot represents a one-minute time-bin from DMT injection, the x-axis reflects the average intensity rating across subjects, and the y-axis indicates the average strength of BW or FW waves across subjects (both correlations p<0.0001). The middle panel shows the correlation coefficients across participants, obtained by correlating the intensity rates and the waves’ amount separately for each time point. Solid lines show when the amount of waves is significantly larger than zero (always for BW waves, few minutes after DMT injections for FW waves – see figure 2A). However, given the limited statistical power (N=12), and proper correction for multiple testing, correlations did not reach significance at any time point. The last panel shows the correlation coefficients between the visual imagery specific ratings provided at the end of the experiment (i.e. Visual Analogue Scale, see methods) and the net amount of waves (measured when both BW and FW were significantly different than zero, i.e. from minutes 2 to 5): for all 20 items in the questionnaire there was a positive trend between the amount of FW waves and the intensity of visual imagery, as confirmed by a Bayesian t-test against zero (BF for FW waves >> 100). We did not observe this effect in the BW waves (BF = 0.41).

## Discussion

In this study we investigated the effects of the classic psychedelic drug DMT on cortical spatio-temporal dynamics typically described as travelling waves ^22^. We analyzed EEG signals recorded from a pool of participants who kept their eyes closed while receiving drug. Results revealed that compared with consistent eyes-closed conditions under placebo, eyes-closed DMT is associated with striking changes in cortical dynamics, which are remarkably similar to those observed during actual eyes-open visual stimulation^16,17^. Specifically, we observed a reduction in BW waves, and increase in FW ones, as well as an overall decrease in α band (8-12Hz) oscillatory frequencies ^7^. Moreover, increases in the amount of FW waves correlated positively with real-time ratings of the subjective intensity of the drug experience as well as post-hoc ratings of visual imagery, suggesting a clear relationship between travelling waves and conscious experience.

### Relation to previous findings

Initiated by the discovery of mescaline, and catalyzed by the discovery of LSD, Western medicine has explored the scientific value and therapeutic potential of psychedelic compounds for over a century ^23–25^. DMT has been evoking particular interest in recent decades however, with new studies into its basic pharmacology ^26^, endogenous function ^27^ as well its effects on cortical activity in rats ^28,29^ and humans ^10,30,31^. There has been a surprising dearth of resting-state human neuroimaging studies of pure DMT ^7,32^ which, given its profound and basic effects on conscious awareness, could be viewed as a scientific oversight.

Previous work involving ayahuasca and BOLD fMRI found increased visual cortex BOLD signal under the drug vs placebo while participants engaged in an eyes-closed imagery task – a result that was interpreted as consistent with the ‘visionary’ effects of ayahuasca ^31^. Despite some initial debate ^33^, it is now generally accepted that occipital visual regions become activated during visual imagery ^34,35^. Placing these findings into the context of previous work demonstrating increased FW travelling waves during direct visual perception ^16,17^, our present findings of increased FW waves under DMT correlating with visionary experiences lend significant support to the notion that DMT/ayahuasca and perhaps other psychedelics, engage the visual apparatus in a fashion that is consistent with actual exogenously driven visual perception. Future work could extend this insight to other apparently endogenous generated visionary experiences such as occur during dreaming and hallucinatory states. We would hypothesize a consistent favoring of FW waves during these dates. If consistent mechanisms were also found to underpin hallucinatory experiences in other sensory modalities – such as the auditory one, a basic principle underlying sensory hallucinations might be established.

### Pharmacological considerations

As a classic psychedelic drug, DMT’s signature psychological effects are likely mediated by stimulation of the serotonin 2A (5-HT2A) receptor subtype. As with all other classic psychedelics ^2^ the 5-HT2A receptor has been found to be essential for the full signature psychological and brain effects of Ayahuasca ^10^. In addition to its role in mediating altered perceptual experiences under psychedelics, The 5-HT2A receptor has also been linked to visual hallucinations in neurological disorders, with a 5-HT2A receptor inverse agonist having been licensed for hallucinations and delusions in Parkinson’s disease with additional evidence for its efficacy in reducing consistent symptoms in Alzheimer’s disease^36^. Until recently, a systems level mechanistic account of the role of 5-HT2A receptor agonism in visionary or hallucinatory experiences has however been lacking.

### Predictive coding and psychedelics

There is a wealth of evidence that predictive mechanisms play a fundamental role in cognitive and perceptual processing ^37,38^ and our understanding of the functional architecture underlying such processing is continually being updated ^16,39^. According to predictive coding ^40^, the brain strives to be a model of its environment. More specifically, based on the assumption that the cortex is a hierarchical system – message passing from higher cortical levels is proposed to encode predictions about the activity of lower levels. This mechanism is interrupted when predictions are contradicted by the lower-level activity (‘prediction error’) – in which case, information passes up the cortical hierarchy where it can update predictions. Predictive coding has recently served as a guiding framework for explaining the psychological and functional brain effects of psychedelic compounds ^20,41^. According to one model ^20^, psychedelics decrease the precision weighting of top-down priors, thereby liberating bottom-up information flow. Various aspects of the multi-level action of psychedelics are consistent with this model, such as the induction of asynchronous neuronal discharge rates in cortical layer 5 ^42^, reduced alpha oscillations ^11,43^ increased signal complexity ^7,12^ and the breakdown of large-scale intrinsic networks ^43^. Recent empirically supported modelling work has lent support to the assumptions that top-down predictions and bottom-up prediction-errors are encoded in the direction of propagation of cortical travelling waves ^16^. This discovery opened-up a tantalizing opportunity for testing assumptions both about the nature of travelling waves and how they should be modulated by psychedelics ^20^. That prior assumptions were so emphatically endorsed by the data, including how propagation-shifts related to subjective experience, enables us to make strong inferences about both the functional relevance of cortical travelling waves and brain action of psychedelics. Future studies can now be performed to examine how these assumptions translate to other phenomena such as non-drug induce visionary and hallucinatory states.

### Conclusion

These are the first EEG data on the effects of DMT on human resting state brain activity. In line with prior hypothesis, clear evidence was found of a shift in cortical travelling waves away from the normal basal predominance of backward waves and towards the predominance of forward waves - remarkably similar to what has been observed during eyes-open visual stimulation. Moreover, the increases in forward waves correlated positively with both the general intensity of DMT’s subjective effects, as well as its more specific effects on eyes-closed visual imagery. These findings have specific and broad implications: for the brain mechanisms underlying the DMT/psychedelic state as well as conscious visual perception more broadly.

## Methods

### Participants and experimental procedure

In this study we analyzed a dataset presented in a previous publication ^7^, to address a very different scientific question using another analytical approach. Consequently, the information reported in this and the next paragraphs overlaps with the previous study (to which we refer the reader for additional details). Thirteen participants took part in this study (6 females, age 34.4 ± 9.1 SD). They all provided written informed consent, and the study was approved by the National Research Ethics (NRES) Committee London – Brent and the Health Research Authority. The study was conducted in line with the Declaration of Helsinki and the National Health Service Research Governance Framework.

Participants were carefully screened before joining the experiments. A medical doctor conducted physical examination, electrocardiogram, blood pressure and routine blood tests. A successful psychiatric interview was necessary to join the experiment. Other exclusion criteria were 1) under 18 years of age, 2) having no previous experience with psychedelic drugs, 3) history of diagnosed psychiatric illnesses, 4) excessive use of alcohol (more than 40 units per week). The day before the experiment a urine and pregnancy test (when applicable) were performed, together with a breathalyzer test. Participants attended 2 sessions, in the first one, they received placebo, while DMT was administered in the second session. Given the lack of human data with DMT, progressively increasing doses were provided to different participants (4 different doses were used: 7mg, 14mg, 18mg and 20mg, to 3, 4, 1 and 5 successive participants respectively). EEG signals were recorded before and up to 20 minutes after drug delivery. Participants rested in a semi-supine position with their eyes closed during the duration of the whole experiment. The eyes closed instruction was confirmed by visual inspection of the participants during dosing. At each minute, participants provided an intensity rating, while blood samples were taken at given time-points (the same for placebo and DMT conditions) via a cannula inserted in the participants’ arm. One day after the DMT session, participants reported their subjective experience completing an interview composed of several questionnaires (see ^7^ for details). In this study we focused on the Visual Analogue Scale values.

### EEG preprocessing

EEG signals were recorded using a 32-channels Brainproduct EEG system sampling at 1000Hz. A high-pass filter at 0.1Hz and an anti-aliasing low-pass filter at 450Hz were applied before applying a band-pass filter at 1-45Hz. Epochs with artifacts were manually removed upon visual inspection. Independent-component analysis (ICA) was performed and components corresponding to eye-movement and cardiac-related artifacts were removed from the EEG signal. The data were re-referenced to the average of all electrodes. All the preprocessing was carried out using the Fieldtrip toolbox ^44^, while the following analysis were run using custom scripts in MATLAB.

### Waves quantification

We epoched the preprocessed EEG signals in 1 second windows, sliding with a step of 500ms (see figure 1). For each time window, we then arranged a 2D time-electrode map composed of 5 electrodes (i.e. Oz, POz, Pz, Cz, FCz). From each map we computed the 2D Fast Fourier Transform (2DFFT -fig.1), from which we extracted the maximum value in the upper and lower quadrants, representing respectively the power of forward (FW) and backward (BW) waves. We also performed the same procedure 100 times after having randomized the electrodes’ order: the surrogate 2D-FFT spectrum has the same temporal frequency content overall, but the spatial information is disrupted, and the information about the wave directionality is lost. In such a manner we obtained the null or surrogate measures, namely FWss and BWss, whose values are the average over the 100 repetitions (see figure 1). Eventually, we computed the actual amount of waves in decibel (dB), considering the log-ratio between the actual and the surrogate values:

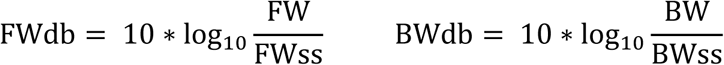

It is worth noting that this value represents the amount of significant waves against the null distribution, that is against the hypothesis of having no FW or BW waves.

### Statistical analysis

All the analyses regarding the EEG signals were performed in Matlab. Bayesian analyses were run in JASP ^45^. We ran separate Bayesian ANOVA for FW and BW conditions, and we considered as factors the time of injection (pre-post, see figure 2A) and drug condition (DMT vs Placebo). Subjects were considered to account for random factors. Regarding the minute-by-minute analysis (figure 2A, right panels) we performed a t-test at each time-bin against zero, and we corrected all the p-values according to the False Discovery Rate ^46^.

## Notes

### Competing Interest Statement

The authors have declared no competing interest.

